# Distinct Myogenic Stages Recapitulate Transcriptomic Networks of Skeletal Muscle in COPD Cachexia

**DOI:** 10.64898/2025.12.03.692216

**Authors:** Hanna Sothers, Bitota Lukusa-Sawalena, Kaleen M Lavin, Wenxia Ma, Joe W Chiles, Richard Casaburi, Rakesh Patel, J Michael Wells, Mohamed Kazamel, Harry B Rossiter, Anna Thalacker-Mercer, Merry-Lynn N McDonald

## Abstract

**Background:** Cachexia is an extrapulmonary manifestation of chronic obstructive pulmonary disease (COPD) characterized by weight loss and muscle wasting. Transcriptomic analysis of skeletal muscle may provide insight to COPD-cachexia-relevant dysregulation, including at the mitochondrial level, where dysfunction is a hallmark of muscle wasting. As muscle biopsies are invasive and yield finite tissue, human muscle derived cultures (HMDCs) can expand limited biopsy material for studying skeletal muscle dysregulation. However, utility of such models depends on their recapitulation of bulk skeletal muscle transcriptomic signatures. To address this, we tested whether COPD and COPD-cachexia dysregulated transcriptional signatures in bulk skeletal muscle are preserved across stages of early differentiation.

**Methods:** *Vastus lateralis* biopsies were collected from 15 (7M/8F, 64±9 years) participants; COPD n=6, COPD cachexia n=4, and n=5 age-matched controls. Cachexia was defined as weight loss coupled with reduced muscle strength, fatigue, anorexia, low muscle mass and/or systemic inflammation. Satellite cells were isolated and differentiated into HDMCs (myoblasts, myocytes, and myotubes). Transcriptomics data was generated from bulk skeletal muscle and HDMCs. Differential expression analysis identified transcripts significantly dysregulated (p<0.05) in COPD and/or COPD-cachexia. Weighted gene co-expression network analysis (WGCNA) preformed at the whole transcriptome and mitochondrial transcriptome levels, identified modules of co-expressed genes. Modules were tested for correlation with clinical traits and preservation between bulk tissue and HDMCs (Z-summary >2). Gene set enrichment analysis was performed for all modules.

**Results:** 660 genes were significantly differentially expressed between COPD and control bulk skeletal muscle. The top upregulated and downregulated genes were *IL21R-AS1* (Log2-Fold-Change [L2FC]=5.3, p=4.2×10^-5^) and top downregulated *POU5F1B* (L2FC=-5.8, p=3.1×10^-2^), respectively. Among the 492 genes significantly differentially expressed between COPD and COPD-cachexia bulk skeletal muscle. The top upregulated and downregulated genes were *LINC02274* (L2FC=5.6, p=3.6×10^-5^) and *ZFY* (L2FC=-6.1, p=9.7×10^-3^) respectively. Modules 1, 9, A, B, D and H correlated with cachexia-relevant traits, and Modules 3, 7, G, and I correlated most strongly with COPD severity. Most modules (1, 2, 4, 7, 8, 9, A, B, H) were preserved across all states; Modules D and G were preserved in myoblasts only, and Module I in myotubes only. Preserved modules were enriched for contractile, inflammatory, oxidative phosphorylation, and fatty acid metabolism pathways.

**Conclusions:** Co-expression modules linked to COPD and COPD-cachexia in bulk skeletal muscle were broadly preserved across all differentiation stages, with myoblast and myotubes most completely recapitulating disease-relevant transcriptional signatures. These findings support HMDCs, as tractable in vitro models preserving contractile, inflammatory, and mitochondrial signatures of COPD-associated muscle dysfunction.

## INTRODUCTION

Chronic obstructive pulmonary disease (COPD) is defined by impairments in pulmonary function and physiology[1]. However, skeletal muscle wasting substantially contributes to limitations in daily activity[2], morbidity, hospitalization, treatment failure and mortality in COPD[3–5]. Important gaps remain in our understanding of the transcriptional and mitochondrial dysregulation underlying muscle wasting in COPD.

Prior research investigating transcriptional dysregulation of skeletal muscle in COPD has largely been restricted to differences between participants with or without COPD rather than between COPD patients with or without cachexia [6, 7]. Yet the muscle transcriptome in COPD cachexia likely differs from that of COPD without cachexia, and this cachexia-specific signature remains poorly characterized. Nonetheless, these studies have revealed dysregulation in pathways involved in oxidative metabolism, extracellular-matrix remodeling, inflammatory signaling, and cell-cycle/repair programs[6, 7]. Specifically, genome-wide profiling of the *vastus lateralis* in COPD demonstrates coordinated dysregulation of mitochondrial and structural pathways. Rabinovich *et al.* identified altered oxidative and energy-metabolism transcripts in patients with reduced fat-free mass index, while Willis-Owen *et al.* reported network-level perturbations involving mitochondrial enzymes and extracellular matrix (ECM) hubs. More recently, our group showed differential co-expression of myofiber transcripts and increased cell-cycle arrest markers (e.g., *CDKN1A*), linking transcriptional remodeling to fiber-type alterations in COPD[8]. Collectively, these patterns point to coordinated disturbances of mitochondrial and structural pathways in skeletal muscle[6, 9–11].

Previous studies of COPD skeletal muscle dysfunction repeatedly implicate mitochondrial pathways, particularly in locomotor muscles such as the quadriceps (*vastus lateralis)*, where impaired bioenergetics and oxidative capacity are well documented[12–14]. Mitochondria are central to skeletal muscle bioenergetics, driving ATP production through oxidative phosphorylation and regulating reactive oxygen species (ROS) and quality-control pathways critical for muscle endurance and repair[15]. In individuals with COPD, skeletal muscle exhibits reduced mitochondrial density and biogenesis, impaired respiratory capacity, imbalanced mitochondrial quality control and elevated oxidative stress, collectively contributing to muscle atrophy, weakness, and diminished independence[16]. This collective maladaptation suggests mitochondrial dysfunction is a central feature of COPD related muscle pathology.

Skeletal muscle transcriptomics can capture COPD and/or cachexia-relevant dysregulation; however, skeletal muscle is a heterogeneous tissue in constant communication with resident and infiltrating cell types. Bulk tissue profiling therefore mixes signals from multiple cell populations, making it difficult to distinguish which dysregulated programs are intrinsic to muscle cells themselves [17–19]. One strategy to alleviate this obstacle is the use of human muscle-derived cultures (HMDC) which consist of satellite cells derived from skeletal muscle lineage: myoblasts and their differentiated progeny, myocytes and myotubes[20, 21]. When activated, skeletal muscle satellite cells differentiate into myoblasts (proliferative), progress to myocytes (early post-mitotic, hybrid state), and finally fuse into myotubes (multinucleated, more oxidative/contractile), a sequence that allows stage-specific interrogation of disease pathways[22]. By enriching for the myogenic compartment, HMDCs isolate muscle-intrinsic transcriptional programs from the confounding heterogeneity of bulk tissue. Although the number of satellite cell are preserved in COPD, their differentiation capacity is impaired suggesting HMDCs capture disease-relevant, muscle-intrinsic dysfunction [23]. While these cultured tissues allow for a convenient *in vitro* model to study tissue dysregulation in a controlled environment, questions remain regarding the extent to which HMDC recapitulate molecular signatures of bulk skeletal muscle in COPD and COPD cachexia. There remains a need to characterize the extent to which stage(s) of *in vitro* differentiation (myoblast, myocyte, or myotube) recapitulate the transcriptional programs of bulk skeletal muscle to evaluate HMDC as tools for mechanistic and translational studies in COPD and COPD cachexia.

To address these gaps, we performed RNA-seq in skeletal muscle from individuals with and without COPD cachexia, in bulk tissue, myoblasts, myocytes, and myotubes. Differential gene expression and weighted gene co-expression network analysis (WGCNA) were used to identify genes and modules associated with COPD and COPD cachexia at the whole and mitochondrial-encoded transcriptome levels. We found that disease-relevant modules were broadly preserved in HMDCs, with myoblast and myotubes most completely recapitulating bulk muscle, supporting HMDCs as a scalable model for studying muscle-intrinsic dysregulation in COPD cachexia..

## METHODS

### Study Design

Participants were recruited to the COPD Cachexia Etiology of Low-Lean (CCELL) Study (NHLBI R01HL153460) from the Birmingham, Alabama metropolitan and surrounding areas. All individuals gave written, informed consent to participate, and all study procedures were approved by the University of Alabama at Birmingham Institutional Review Board. Participants ages 40 to 80 years, with or without COPD, were recruited if willing to undergo study procedures. COPD was diagnosed using post-bronchodilator lung function testing as a forced expiratory volume in one second to forced vital capacity (FEV_1_/FVC) ratio <0.7[24]. FEV_1_ was reported as a percentage of the predicted value (FEV_1_%pred) using race-neutral Global Lung Initiative equation values[25]. GOLD scores were calculated for participants with COPD[26]. COPD and controls with smoking exposure were required to have at least a 10 pack-year smoking history. Participants were excluded if they were pregnant or breastfeeding, had significant ongoing alcohol dependency or drug abuse, had history of anorexia or bleeding disorders, had cancer within the past 7 years, HIV/AIDS, long-term anticoagulant use, history of Methicillin-Resistant Staphylococcus aureus (MRSA) infection, BMI ≥ 35 kg/m², or had a COPD exacerbation within the last three months.

### Clinical Phenotypes of COPD

#### Emphysema

Chest CT scans were acquired at full inspiration using standardized protocols. All CT images underwent automated lung segmentation using in-house algorithms to isolate the lung parenchyma. Emphysema quantifications were performed using low attenuation areas (LAA) based on emphysematous density thresholds[27]. Percentage of lung volume with attenuation values less than-950 Hounsfield units (HU), representing emphysematous areas (LAA-950).

### Presence of COPD Cachexia

Presence of COPD cachexia was determined based on consensus criteria developed by Evans and colleagues[24]. Only participants with COPD were assessed for the presence of cachexia. Cachexia was defined as participant-reported weight loss ≥ 5% total body mass over the past 12 months or ≥ 2% in past 12 months if BMI <20 kg/m^2^, in addition to at least 3 of the 5 following criteria:

#### Anorexia

Anorexia was assessed using the Simplified Nutritional Appetite Questionnaire[28]. Participants with a total score < 14 were considered to have anorexia[29].

#### Abnormal Blood Biochemistry

Serum from a peripheral blood sample was allowed to clot at room temperature for 30-60 min, aliquoted, and immediately frozen until measurement of key circulating factors. Albumin was measured with a Randox (Crumlin, UK) RX Daytona+ Clinical Chemistry Analyzer. Growth and differentiating factor (GDF)-15 was measured in duplicate using an ELISA kit. Interleukin (IL)-6 was measured in duplicate using MesoScale Discovery (Rockville, MD) Human IL-6 kits using chemiluminescence. C-reactive protein (CRP) was measured with a Randox (Crumlin, UK) RX Daytona+ Clinical Chemistry Analyzer. Using automated multichannel hematology analyzers, hemoglobin (Hgb) was quantified spectrophotometrically via the sodium lauryl sulfate (SLS) hemoglobin method. Hematocrit (Hct) was derived from the red blood cell count and mean corpuscular volume using the analyzer’s standard algorithm[30, 31]. For each biomarker, samples that fell above or below the limit of detectability on a given assay were imputed with a conservative value ±0.001 units above or below the threshold, respectively. This was done to retain a value for the sample enabling downstream statistical analysis that relies on a full-rank matrix (e.g., correlations). From these, presence of either anemia, hypoalbuminemia, or increased serum inflammatory markers was used to denote abnormal blood biochemistry as follows: Anemia: Hgb < 12 | 13.6 g/dL or hematocrit (Hct) <37% | 40% (females | males)[32]; serum albumin < 3.5 g/dL[33]; CRP > 5mg/L, IL-6 > 4pg/mL[34], or GDF-15 > 1500 pg/mL..

#### Fat Free Mass Index

A total body scan was performed using the Lunar iDXA (General Electric, Cincinnati, OH). Total lean mass divided by the square of height was used to calculate FFMI, which was dichotomized as normal or low based on thresholds developed by Schols, *et al.*[35] (<15 kg/m^2^ in females, <16 kg/m^2^ in males).

#### Fatigue

Fatigue was assessed using the Fatigue Severity Scale, a nine-question assessment scoring items using a seven-point Likert scale. Participants with an average score ≥4 points were considered to have fatigue, based on ATS guidelines[36–38].

#### Poor Muscle Strength

Hand grip strength (HGS) was measured via dynamometer (Biodex Medical Systems, New York, NY). The best of three attempts was taken as the maximum HGS and used to determine low HGS, defined as <20 kg in females and <30 kg in males[39].

#### Citrate Synthase Activity

From snap frozen bulk skeletal muscle, Citrate Synthase Enzyme activity was measured using the coupled reaction with oxaloacetate, acetyl-CoA, and 5,5-dithiobis-(2,4-nitrobenzoic acid) to provide a surrogate index of mitochondrial volume.

### Muscle Tissue Collection and Processing

*Vastus lateralis* biopsies were obtained under local anesthesia using a Bergström needle with suction[40]. Tissue was cleaned of fat/blood/connective tissue and partitioned for downstream scientific objectives. Approximately 25 mg were set aside for “bulk” RNA extraction, frozen immediately in liquid nitrogen and stored at-80°C until RNA extraction using TRIZol reagent (Ambion, Waltham, MA).

Another 50-75 mg piece for cell culture was set aside for the isolation of human primary myogenic progenitor cells, (HMPCs), as described in detail previously[41]. Briefly, the muscle fiber bundle was preserved in Hibernate-A Buffer (Invitrogen, Waltham, MA) at 4°C for 24-48 hours, minced and washed via gravity with Dulbecco’s PBS (Gibco, Waltham, MA) and then digested using mechanical and enzymatic digestion in low-glucose Dulbecco’s Modified Eagle Medium (Gibco). This solution was passed through a 70-mm cell strainer into 5 mL of a growth medium comprised of Ham’s F12 (Gibco), 20% FBS, 1% penicillin/streptomycin (Corning, Corning, NY), and 5 ng/mL of recombinant human basic fibroblast growth factor (Promega, Madison, WI) and then centrifuged. The pelleted cells were resuspended in 10% DMSO and cryopreserved at-80 °C until isolation via flow cytometry. Primary HMPCs were sorted using fluorescence-activated cell sorting with fluorescence-conjugated antibodies to cell surface antigens specific to HMPCs (CD56 [NCAM, BD Pharmingen, Franklin Lakes, NJ] and CD29 [b1-integrin, BioLegend, San Diego, CA]) and a viability stain (7-Aminoactinomycin D, eBioscience, San Diego, CA).

Passage six HMPCs were used for all listed experiments and were cultured in a 5% CO_2_ atmosphere at 37°C on collagen-coated plates (Type I, rat tail, Corning) in primary growth media containing in Promocell Human Skeletal Muscle Growth Media + supplement, GlutaMax, Invitrogen Antibiotic-Antimycotic, and 20% FBS. Media was changed every 48 hours until the desired cell count was reached, at which point cells were switched to differentiation media containing Promocell Human Skeletal Muscle Differentiation Media + supplement, GlutaMax, Invitrogen Antibiotic-Antimycotic, and 2% horse serum. From each participant, RNA was harvested at three time points: day 0 (myoblast stage), day 3 (myocyte stage), and day 6 (myotube stage). RNA extraction was performed using the Omega EZNA Total RNA Kit I (Omega, Norcross, GA) and quantified spectrophotometrically.

### Generation of RNA-Seq Data and Quality Control

RNA from bulk skeletal muscle homogenate as well as derived myoblasts, myocytes, and myotubes were assessed for concentration via NanoDrop (504 ± 265 ng/µL) and quality via Agilent bioanalyzer (RIN: 9.2 ± 1.2; 28S/18S ratio: 1.4 ± 0.8) from 13 CCELL study participants. Samples were sequenced on the Illumina NovaSeq 6000 using paired end sequencing. Resulting FASTQs were assessed for quality using FastQC[42] and Illumina universal adapter (sequence: AGATCGGAAGAG) were removed using cutadapt[43]. Resulting reads below 15bp were dropped from analysis. Samples were aligned using STAR aligner version 2.7.3 to GRCh38 in NCBI with a sjdb overhang setting of 100 bp. Samples with aligned read totals below 10 million (n=2) were interrogated using principal components analysis to identify whether they were extreme data points on a global gene expression scale, in which case they were dropped from analysis (n=0). Mean ± SD uniquely mapped read totals were 31.7 million ± 8.4 million.

### Descriptive Statistics

A t-test was used to assess the difference in FEV1%pred between all COPD cases and controls. For comparisons between the non-COPD, COPD, and COPD cachexia groups, a Kruskal-Wallis test with Dunn correction was used for continuous variables and chi-square was used for categorical variables

### Differential Gene Expression Analysis

DGE analysis was performed using DESeq2 (v1.44.0)[44] in R (v4.4.0). A raw count matrix was generated from bulk skeletal muscle RNA-seq data. Gene expression was filtered to retain only those with expression >0 in at least 25% of the samples (i.e., ∼3 samples). Count data were normalized using DESeq2’s median of ratios method to account for library size and RNA composition differences. Variance-stabilizing transformation (VST) was applied for visualization and downstream correlation analyses. Genes with |*log2 fold change (L2FC)*| > 1 and unadjusted p < 0.05 were considered significantly differentially expressed. Volcano plots were generated using R packages colorspace (v2.1.2), ggplot (v), and ggpub (v0.6.2).

### Weighted Gene Correlation Network Analysis

#### Network Construction – Whole Transcriptome

Transcriptional data for each HMDC state and the bulk skeletal muscle were treated independently using an identical workflow to account for differences in the data structure. Gene expression was filtered as stated previously in analysis. VST was applied independently to each data set for normalization, as recommended by WGCNA developers [45].

WGCNA (v1.73) was used to construct networks of modules based on pairwise relatedness between genes for bulk skeletal muscle and the three HMDC states. Soft thresholding power was determined independently for each data set prior to the computation. Signed correlation networks were built using a bi-weight mid-correlation with the following parameters: minimum module size to was set to 30, and *mergeCutHeight* to 0.5. In each data set, genes that did not meet the criteria for relatedness to any module were placed in the “grey” category and excluded. A kME > 0.8 was used to identify hub genes [46, 47].

#### Network Construction – Mitochondrial Transcriptome

Similarly, WGCNA was performed at the mitochondrial transcriptome level, using a curated subset of 1,136 mitochondrial and nuclear encoded mitochondrial genes from the Broad Institute MitoCarta 3.0 catalog [48] to further investigate the role of mitochondrial dysfunction in COPD cachexia. To accommodate for the notably smaller gene set, parameters were adjusted to increase sensitivity for detecting smaller, functionally distinct modules. Specifically, deepSplit was set to 3 to allow finer resolution of modules within this restricted gene set, and mergeCutHeight was lowered to 0.25 to avoid prematurely merging biologically distinct mitochondrial processes.

#### Module Preservation

Following network construction, module preservation analysis was performed in the WGCNA package to determine whether network modules were conserved between bulk skeletal muscle and each of the HMDC states. We defined preservation as the retention of co-expression topology for bulk-defined modules in HMDCs quantified by a Z-summary statistic. This statistic is a measure of module similarity[49], where a higher value indicates a higher likelihood than random chance of finding genes together in a module. A Z-Summary score above 2 indicates moderate preservation, and a score above 10 indicates a high degree of preservation. Module preservation was evaluated independent of phenotype.

#### Module-Trait Correlation

Bulk skeletal muscle co-expression modules were assessed for correlation with COPD and cachexia related traits using Pearson correlation (r). Gene-trait correlation and significance was calculated for all significantly correlated traits for each module. P-values were calculated using *corPvalueStudent* from the WGCNA package. Correlation heatmap was also generated using the WGCNA package.

### Gene Set Enrichment Analysis (GSEA)

Gene set enrichment analysis was performed using the enrichR package[50–52]. The following databases were queried: SysMyo, MoTrPAC(2023), Gene Ontology: Biological Processes, Cellular Component, and Molecular Function(2025), MSigDB(2020), KEGG(2021), Reactome(2024), and WikiPathways(2024). Gene sets with an unadjusted p-value < 0.05 or less than 2 overlapping genes were not considered.

## RESULTS

### Participant Demographics

Among the 15 participants with bulk skeletal muscle (n=5 non-COPD controls, n=6 COPD, and n=4 COPD-cachexia cases) included in this study there was no significant differences in age between the three groups (Table 1). Serum CRP was significantly higher in COPD cachexia cases compared to controls. As expected, lung function (FEV_1_%pred) was lower in all participants with COPD compared to the controls (p=0.001, mean FEV_1_%pred: 61.7% and 95.5%, respectively).

**Table 1:** Characteristics of CCELL participants with bulk skeletal muscle samples used in this study.

|  | Control | COPD | COPD Cachexia |
| --- | --- | --- | --- |
| **Demographics** |  |  |  |
| n | 5 (3M, 2F) | 6 (4M, 2F) | 4 (0M, 4F) |
| Age | 69 (13) | 65 (14) | 58 (3) |
| Ever Smoker | 3 (60%) | 6 (100%) | 4 (100%) |
| **Physical Characteristics** |  |  |  |
| % Emphysema | 2.7 (1.2) | 5.2 (7.1) | 1.5 (0.5) |
| Anorexia n(%) | 1 (20%) | 2 (33%) | 3 (75%) |
| Fatigue n(%) | 1 (20%) | 2 (33%) | 4 (100%) |
| BMI (kg/m<sup>2</sup>) | 29 (2) | 28 (6) | 34 (4) |
| Low FFMI | 2 (40%) | 2 (33%) | 3 (75%) |
| Low HGS | 0 (0%) | 0 (0%) | 2 (50%) |
| **Blood Biomarkers** |  |  |  |
| Albumin (g/dL) | 44 (1) | 44 (2) | 43 (2) |
| Hct (g/dL) | 41 (7) | 46 (4) | 39 (5) |
| Hgb (g/dL) | 14 (2) | 15 (1) | 13 (2) |
| CRP (mg/L) <sup>*,††</sup> | 3.1 (1.8) | 6.3 (5.1) | 11 (6.3) |
| GDF-15 (pg/mL) | 1100 (480) | 1300 (820) | 1400 (200) |
| IL-6 (pg/mL) | 0.92 (0.32) | 1.7 (1.3) | 1.4 (0.2) |

Note: Data presented as median (interquartile range)
Abbreviations: F: female, M: male, FEV1: Force Expiratory Volume in 1 second, BMI: body mass index, HGS: handgrip strength, FFMI: fat-free mass index, Hct: hematocrit, Hgb: hemoglobin, CRP: C-Reactive Protein, GDF-15: Growth Determination Factor 15, IL-6: Interleukin 6
<sup>\*</sup>denotes post-hoc p value < 0.05 for COPD vs Controls, and <sup>††</sup> p < 0.01 COPD-Cachexia vs Controls.

### COPD and COPD Cachexia Significantly Differentially Expressed in Bulk Muscle Biopsy Samples

A total of 666 genes were significantly differentially expressed (228 upregulated genes, 438 genes downregulated) in the bulk skeletal muscle of all COPD participants compared to controls (Figure 2a; Table S1). The top upregulated gene was *IL21R-AS1* (Interleukin 21 Receptor Antisense RNA 1) (L2FC=5.3, p=4.2×10^-5^) and top downregulated *POU5F1B* (POU class 5 homeobox 1B) (L2FC=-5.8, p=3.1×10^-2^). A total of 497 genes were significantly differentially expressed (112 genes upregulated, 385 genes downregulated) in bulk skeletal muscle from COPD participants with cachexia versus those without (Figure 2b; Table S2). The top upregulated gene was *LINC02274* (Long intergenic non-protein coding RNA 2274)(L2FC=5.6, p=3.6×10^-5^) and top downregulated *ZFY* (zinc finger protein Y-linked) (L2FC=-6.1, p=9.7×10^-3^).

**Fig. 1.**
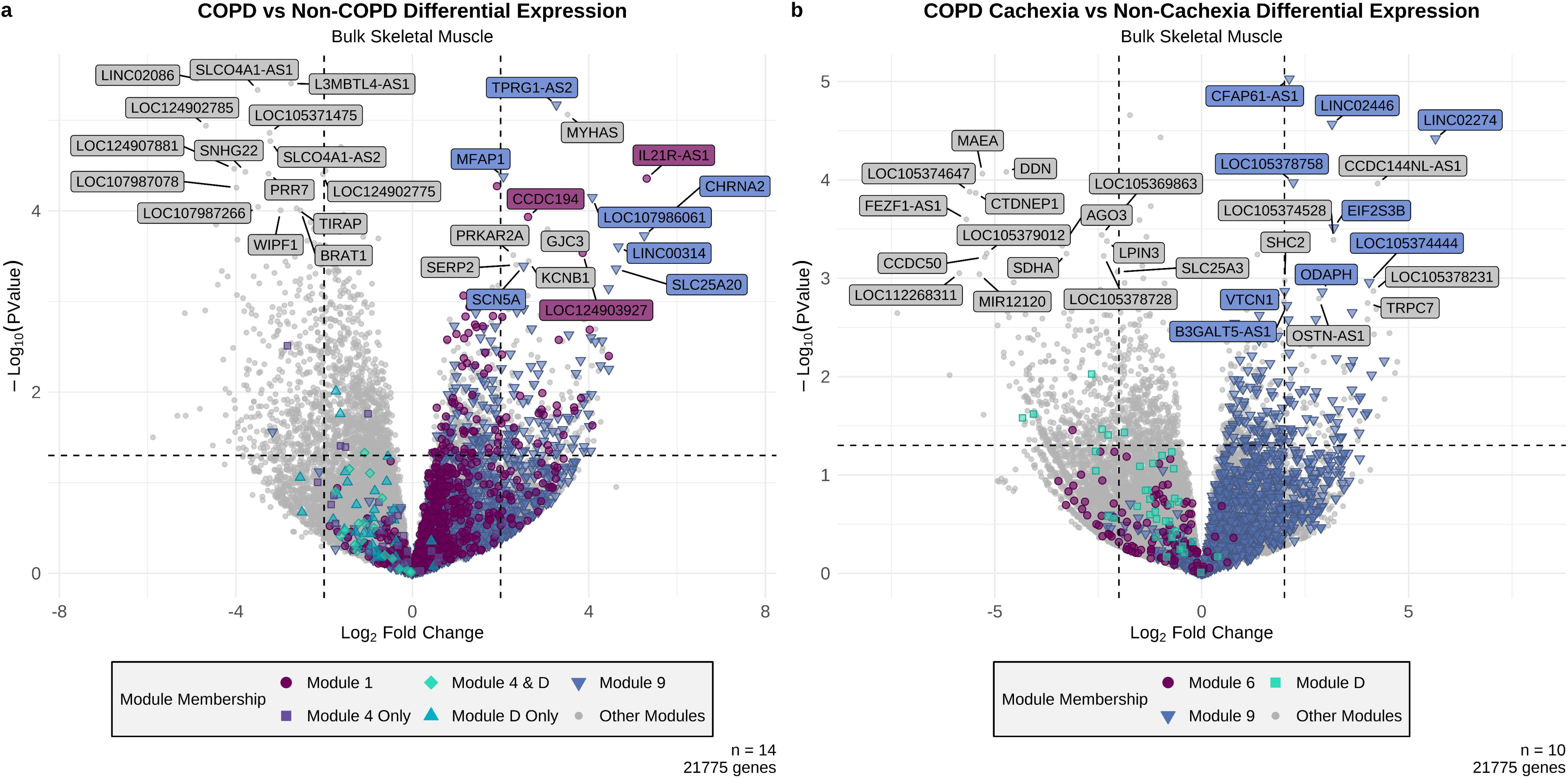
Graphical Abstract. Schematic of *vastus lateralis* biopsies from older adults with and without COPD with and without cachexia provided bulk muscle for RNA-seq and satellite cells that were differentiated *in vitro* (D0 myoblasts → D3 myocytes → D6 myotubes). We used differential gene expression (DGE) analysis to profile tissue-level changes, then applied weighted gene co-expression network analysis (WGCNA) to build modules, test module preservation from tissue to culture stages, and relate modules to clinical traits. Gene set enrichment analysis (GSEA) annotated preserved networks, highlighting disease-relevant pathways and the culture stages that best recapitulate mature muscle programs. Figure was created in BioRender.

**Fig. 2.**
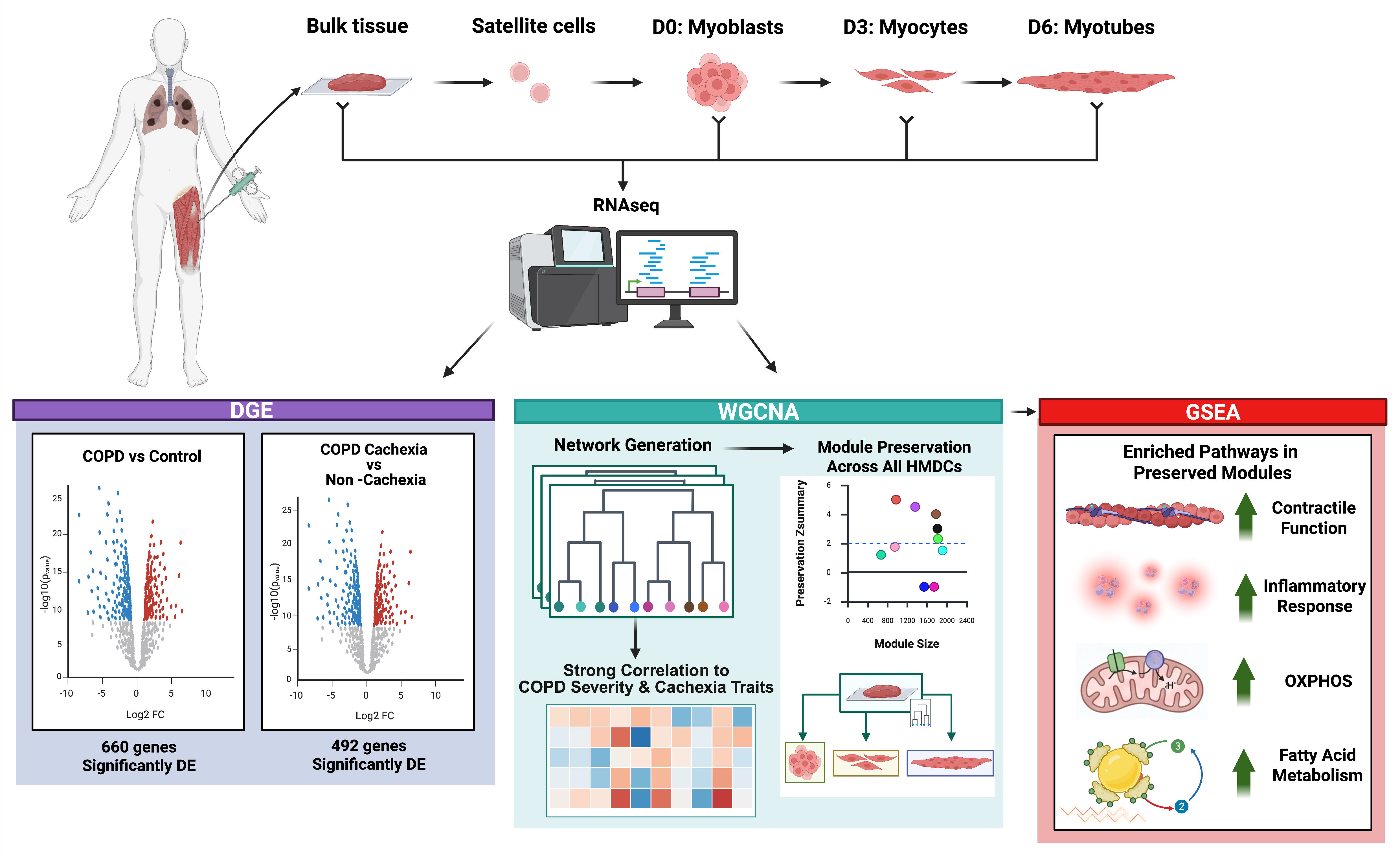
Differential gene expression analysis of All COPD vs Control and COPD vs COPD Cachexia. Volcano plot of differentially expressed genes in (a) All COPD individuals compared to the controls and (b) COPD individuals compared to those without cachexia. Genes with a p-value <0.05 & |Log2FoldChange| > 1 were considered significantly differentially expressed. The top 15 significantly up-and down-regulated genes are labeled. Points are colored by module membership for modules significantly correlated to COPD (a) or COPD Cachexia (b). Top 15 upregulated and top 15 downregulated genes that met threshold of |Log2FoldChange| > 1 after p-value filter of <0.05 were labeled and colored genes are within correlating modules. Plots were created in R using the ggplot2 package.

### Assessment of Preservation of Bulk Muscle Biopsy Co-Expression Modules in Derived Myoblasts, Myocytes, and Myotubes

Most modules were preserved in at least two of the three HMDC differentiation states. Among the whole transcriptome modules, Modules 1, 2, 4, 7, 8, and 9 were preserved across all three states while Module 3 was preserved in only myoblasts and myotubes, and Module 6 preserved in only myocytes and myotubes (Figure S1). Among the mitochondrial transcriptome modules, Modules A, B, and H were preserved in all three states. Modules D and G were preserved in myoblasts only, while Module I was preserved in myotubes only.

### COPD Severity and Cachexia-Related Correlations in the Whole Transcriptome Network

WGCNA of the bulk skeletal muscle generated 10 modules from the whole transcriptome (designated modules 1-10) and 10 modules from the mitochondrial transcriptome (designated modules A-I) (Tables S6-23). Module preservation scores between bulk skeletal muscle and each of the HMDC states are shown in Figure S1. Among the whole transcriptome modules, Module 1 showed a significant negative correlation with FFMI (r = −0.6, p =0.03) and was enriched for amino acid transport processes, specifically L-arginine transmembrane transport, demonstrating that lower lean mass index is associated with higher expression of this module (Figure 3; Table S24).. Module 3 showed the strongest trait associations, with a significant positive correlation with COPD GOLD stage (r = 0.8, p =5×10⁻⁴) and a significant negative correlation with FEV1% predicted (r =−0.7, p =6×10^-3^), consistent with this module tracking with disease severity. Module 3 was strongly enriched for genes involved in muscle contraction, sarcomere organization, myofibril assembly, and actin-myosin filament sliding (Table S26). Module 6 showed a significant positive correlation with hemoglobin (r = 0.7, p =0.01) with enrichment in ATP biosynthesis and extracellular matrix disassembly (Table S29). Module 7 was positively correlated with COPD GOLD stage (r = 0.7, p =4×10^-3^) and enriched for stress response, genome maintenance, and intracellular trafficking, among other pathways (Table S30). Module 8 showed a significant negative correlation with CRP (r = −0.5, p =0.042), This is consistent with enrichment of genes involved in extracellular matrix assembly, *TGF-β* signaling, and cytoskeletal organization (Table S31). In contrast. Module 9 demonstrated a coordinated inflammatory signature, with significant negative correlation with FFMI (r = −0.6, p =0.02) and significant positive correlations with COPD cachexia (r = 0.6, p =0.02) and CRP (r = 0.8, p =4×10⁻⁴), suggesting a role in inflammation-driven muscle wasting indicating an inverse relationship with systemic inflammation relative to Module 8. Genes in this module were enriched for cardiac and muscle regulatory processes (Table S32)

**Fig. 3.**
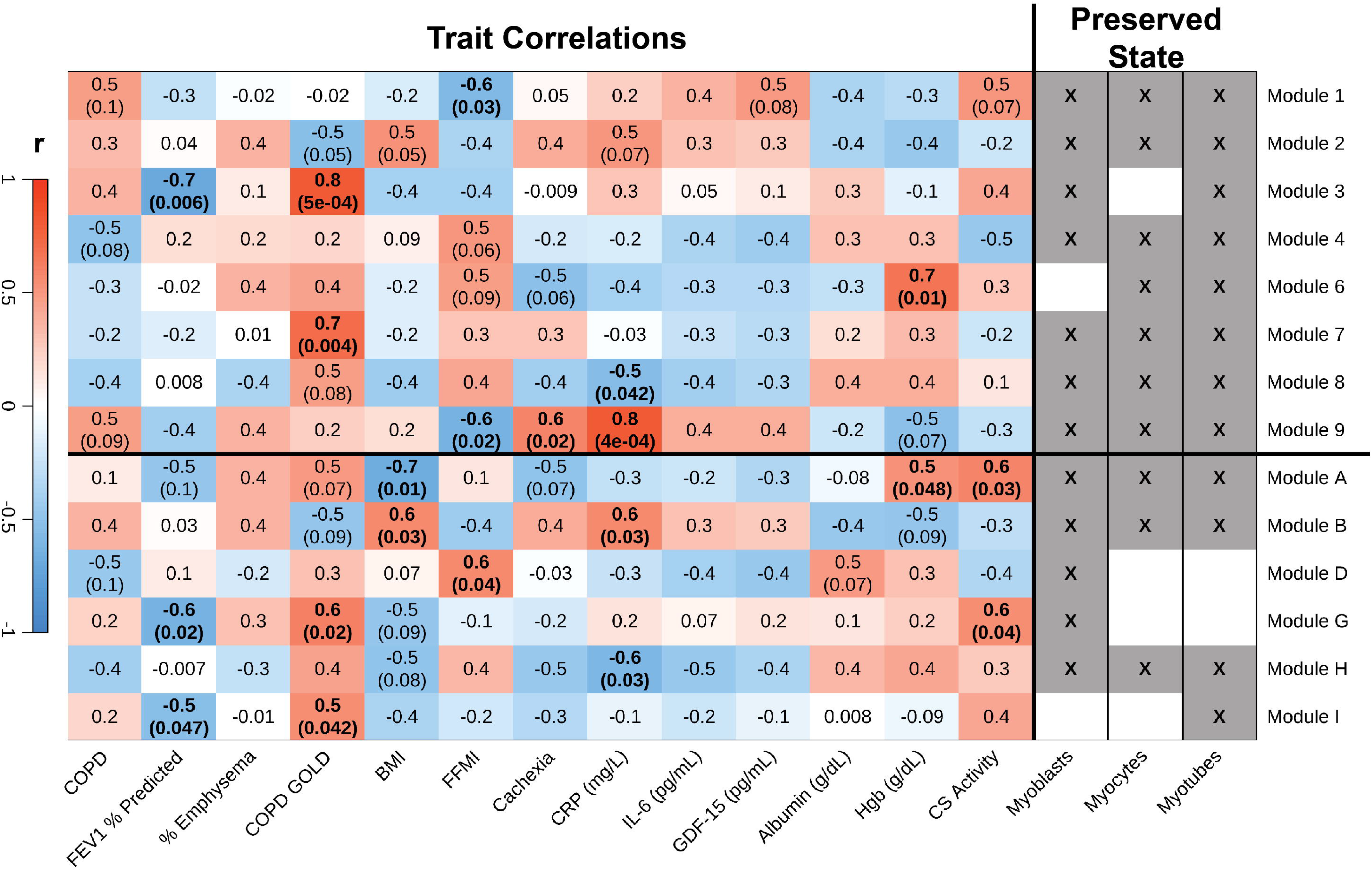
Module-Trait Correlation of the Preserved Bulk Tissue Modules and COPD/Cachexia-related Traits. Module-Trait Correlation of the Preserved Bulk Tissue Modules and COPD/Cachexia-related Traits. Columns 1-13 show correlation (Pearson’s) of selected traits to the module eigengenes. Columns 14-16 show which HMDC state each mature tissue module was preserved. For significant correlations (p <0.05), the p-value is listed below r in parentheses and the text is bolded. P-value is placed underneath in parentheses if p <0.05 and text is bolded. Modules preserved in myoblasts, myocytes, or myotubes are indicated with an X mark. Image was created in R using the WGCNA package.

### Mitochondrial Network Reflects Diverse Bioenergetic and Regulatory Programs

Within the mitochondrial transcriptome modules, Module A showed significant negative correlation with BMI (r = −0.7, p = 0.01) and positive correlations with hemoglobin (r = 0.5, p = 0.048) and CS activity (r = 0.6, p = 0.03). This module is enriched for negative regulation of ferroptosis, mitochondrial respirasome assembly, and protein localization to the mitochondrion (Table S33). Module B was positively correlated with both BMI (r = 0.6, p = 0.03) and CRP (r = 0.6, p = 0.03) with strong enrichment for mitochondrial translation, protein import into the mitochondrial matrix, and carnitine metabolic processes (Table S34). Module D showed a significant positive correlation with FFMI (r = 0.6, p = 0.04) and enrichment for mitochondrial ribosome assembly, and phosphatidylglycerol biosynthesis (Table S36), consistent with a role in lean mass maintenance. Module G showed significant associations with FEV1% predicted (r = −0.6, p = 0.02), COPD GOLD stage (r = 0.6, p = 0.02), and CS activity (r = 0.6, p = 0.04), positioning it as a module tracking both airflow limitation and mitochondrial oxidative capacity. Module G was enriched for calcium import regulation, mitochondrial electron transport and ketone body metabolism (Table S39). Module H was significantly negatively correlated with CRP (r = −0.6, p = 0.03). Genes in this module were strongly enriched proton motive force-driven ATP synthesis, oxidative phosphorylation, cellular respiration, and ATP biosynthesis (Table S40). Module I was negatively correlated with FEV1 % predicted (r = −0.5, p = 0.0.047) and positively correlated with COPD GOLD stage (r = 0.5, p = 0.042), indicating a complex relationship with disease severity and body composition. Genes in this module showed enriched for fatty acid beta-oxidation, mitochondrial electron transport from ubiquinol to cytochrome c, and protein repair (Table S41).

## DISCUSSION

Mechanistic study of skeletal muscle in COPD cachexia is confined by the limited yield of muscle biopsies, restricting the scale at which disease-relevant dysregulation can be investigated. A human-derived culture model that recapitulates bulk skeletal muscle transcriptional networks would help overcome this barrier. Here, by comparing bulk skeletal muscle transcriptomics with HMDCs, we found that several co-expression modules identified in bulk skeletal muscle are preserved across myoblast, myocyte, and myotube stages supporting HMDCs as biologically relevant platforms for studying COPD cachexia. We discovered that several key co-expression modules identified in bulk skeletal muscle are preserved in myoblasts, myocytes and myotubes, supporting the use of these stages as biologically relevant platforms for studying degeneration and repair pathway dysregulation in COPD cachexia.

The pattern of whole transcriptome module-trait correlations reveals a coherent biological narrative connecting disease severity, body composition, and systemic inflammation in skeletal muscle of individuals with COPD. Module 3 showed the most intermodular connectivity, enriched for sarcomere organization and contractile mechanisms, with the most severe COPD GOLD stages and lowest FEV1 values suggests that progressive airflow obstruction is accompanied by coordinated suppression of the muscle contractile transcriptome, which is consistent with the well-documented fiber-type cross-sectionally and contractile protein loss in advanced COPD [11, 53]. The opposing correlations of Modules 8 and 9 with CRP are particularly informative: the positive association of Module 9 with both COPD cachexia and CRP alongside its negative association with FFMI positions it as a potential transcriptional hub for inflammation-driven wasting. Meanwhile, This divergence raises the question of whether skeletal muscle in COPD is simultaneously engaged in inflammatory catabolism and structural remodeling, and whether these processes compete. Because CRP reflects systemic rather than muscle-specific inflammation, distinguishing muscle-intrinsic inflammatory signaling from a systemic response will require direct measurement of local inflammatory mediators.. The negative correlation of Module 1 with FFMI, combined with enrichment for amino acid transport, is consistent with the known impairment of protein anabolic signaling in cachectic muscle, where reduced amino acid uptake capacity may compound the effects of systemic inflammation on net protein balance [54, 55]. Our co-expression modules retain this established biology supports the validity of the network framework, providing confidence that preserved modules without prior characterization reflect genuine muscle programs rather than technical artifacts.

Within the mitochondrial transcriptome, the pattern of module-trait associations points to distinct layers of mitochondrial dysfunction at different levels of the Oxidative phosphorylation (OXPHOS) hierarchy. At the ATP synthesis and oxidative capacity level Module H, enriched for the core proton motive force-driven ATP synthesis machinery and negatively correlated with CRP, suggests that higher expression of this OXPHOS module is associated with lower systemic inflammation, potentially reflecting a protective bioenergetic capacity against inflammation-driven muscle catabolism [13, 56]. Consistent with this, Module G’s association with both airflow obstruction and CS activity positions it at the intersection of systemic disease severity and mitochondrial oxidative capacity, consistent with evidence that CS activity declines with COPD progression[57]. In the context of substrate oxidation Module I, negatively correlated with FEV1% predicted and positively correlated with COPD GOLD stage, suggests that fatty acid beta-oxidation and electron transport programs track with disease severity rather than the cachectic phenotype specifically. Module A’s negative correlation with BMI and positive correlations with hemoglobin and CS activity may reflect a biologically plausible link between mitochondrial respirasome assembly, ferroptosis regulation and iron-dependent processes relevant to both oxygen-carrying capacity and mitochondrial oxidative function, given the established role of iron in both processes [58, 59]. In contrast, Module B, positively correlated with both BMI and CRP, suggests that mitochondrial translation and protein import programs are elevated alongside systemic inflammatory burden, potentially reflecting a stress-responsive upregulation of mitochondrial biogenesis under inflammatory conditions. This is consistent with evidence that mitochondrial protein import and biogenesis programs can be modulated by inflammatory signaling in skeletal muscle [60, 61]. Together, these module associations suggest that cachexia in COPD involves not only reduced OXPHOS output but also disrupted substrate oxidation and potentially maladaptive compensatory translation, demonstrating a multi-layered pattern of mitochondrial transcriptional dysfunction that may be more comprehensively captured by the targeted mitochondrial network approach than by whole transcriptome analysis alone. This approach provides sufficient representation and finer resolution of mitochondrial signatures that may be dispersed across modules in the whole transcriptome network. Given the central role of mitochondrial dysfunction in COPD muscle, this resolution offers a framework for targeted mechanic follow-up.

The preservation patterns across HMDC differentiation states carry important implications for the selection of appropriate in vitro models. Because myotubes are differentiated and closest to multinucleated, oxidative phenotype of mature muscle fibers, we anticipated that later differentiation stages would most accurately recapitulate the bioenergetic and contractile programs of bulk muscle [62, 63]. The broad preservation of Modules 1, 2, 4, 7, 8, and 9 across all three HMDC states in the whole transcriptome network, and of Modules A, B, and H in the mitochondrial network, indicates that the core transcriptional programs associated with COPD-relevant traits are robustly maintained across the full differentiation trajectory. The selective preservation of Module 3 in myoblasts and myotubes but not myocytes is particularly notable given its strong association with disease severity. The absence of this contractile program in myocytes likely reflects the transient suppression of structural muscle genes during the proliferative-to-differentiative transition, rather than a true loss of disease relevance[64, 65]. This observation is consistent with the transitional nature of the myocyte state, in which transcripts associated with both proliferation and terminal differentiation are simultaneously active, potentially diluting stable module structure [66]. Module I’s exclusive preservation in myotubes aligns with the biological expectation that fatty acid beta-oxidation programs reach full transcriptional maturity only in terminally differentiated cells, further supporting the myotube as the most appropriate stage for modeling mature bioenergetic phenotypes [67]. Module D and Module G’s restriction to myoblasts suggests that mitochondrial gene expression, ribosome assembly, and calcium-mitochondria coupling programs are most transcriptionally active, proliferating myogenic cells. since cultured myoblasts represent an activated rather than quiescent state and differ substantially from resident satellite cells in vivo, these programs are best interpreted as reflecting activated regenerative activity rather than the behavior of inactive satellite cells in COPD muscle[68, 69].

Interestingly, despite strong preservation across differentiation states, several mitochondrial modules did not show significant association with cachexia or COPD status as a trait, suggesting these core mitochondrial programs are relatively stable at the transcript level in this cohort and COPD cachexia-related mitochondrial dysfunction may be better captured by functional or multi-omics readouts. Therefore, the preserved HMDC models provide a practical platform for targeted mechanistic follow-up of genes in the inflammatory and mitochondrial modules through targeted knockdown, overexpression, or metabolic and inflammatory challenges. Identifying which of these programs are muscle-intrinsic and modifiable could reveal therapeutic targets for catered to skeletal muscle wasting in COPD cachexia. Prior transcriptomic studies of skeletal muscle in COPD and cachexia have consistently reported broad patterns of dysregulation, including abnormal myofiber-type, impaired oxidative metabolism, and enrichment of inflammatory pathways [8, 70, 71]. Here, we extend these observations by showing that key co-expression programs detected in vivo are retained in patient-derived culture stages. The strong preservation observed at the myotube stage aligns with prior work showing that differentiated myotubes most closely approximate in vivo muscle physiology among culture states reinforcing the biological validity of this stage for modeling mature bioenergetics and contractile regulation[63, 72]. In addition, module preservation and trait-correlation patterns indicate that myocytes retain several disease-relevant co-expression programs, including modules associated with systemic inflammation and body composition, suggesting partial recapitulation of COPD cachexia-relevant transcriptional programs at this earlier transitional stage. However, modules most strongly tracking disease severity, including the core contractile module and fatty acid oxidation programs, were not preserved in myocytes, underscoring that the myotube stage remains the most complete model of bulk muscle transcriptional architecture [73].

The single-gene level findings provide additional mechanistic context for the network-level observations. The broad downregulation signal in COPD is consistent with suppression of normal homeostatic muscle programs as a dominant feature of the disease transcriptome [7, 74]. Among the cachexia-specific findings, the upregulation of *SEMA4F* (Semaphorin-4F) and *CXCL13* (Chemokine (C-X-C motif) ligand 13) alongside downregulation of the E3 ubiquitin ligase *MAEA* (macrophage-erythroblast attacher) and ribosomal protein *RPL22* suggests that cachexia adds a distinct layer of immune and semaphorin signaling onto the broader COPD transcriptional background, while simultaneously impairing translational capacity and ubiquitin-mediated protein quality control. The upregulation of *SPX* (spexin), a neuropeptide associated with metabolic regulation and energy homeostasis that promotes myoblast proliferation, in COPD cachexia may reflect a compensatory attempt to couple satellite cell-mediated regeneration with metabolic adaptation in response to ongoing muscle catabolism[75]. Taken together, these gene-level signals are coherent with the module-level observation that cachexia engages both inflammatory/catabolic and translational dysregulation programs on top of the disease-severity-associated contractile and metabolic changes captured in the WGCNA modules.

A major strength was recruitment of a cohort phenotyped for both COPD and cachexia outcomes including collection of a muscle biopsy from the vastus lateralis and subsequent preparation of cell cultures from these specimens. Participants were rigorously characterized for COPD severity and cachexia using state-of-the-art clinical criteria, and samples underwent high-depth RNA sequencing, enhancing sensitivity to detect coordinated transcriptional programs. The combination of DGE analysis and WGCNA across whole-transcriptome and mitochondria-focused datasets provided complementary resolution at the single-gene and network levels. The targeted mitochondrial WGCNA further refined mechanistic insight by isolating co-regulated mitochondrial and nuclear-encoded mitochondrial genes that might otherwise be diluted in whole transcriptome analyses. To our knowledge, this is among the first studies to systematically.evaluate whether co-expression networks derived from bulk skeletal muscle in COPD and COPD cachexia are structurally preserved across HMDC states. Nonetheless, the relatively small sample size and lack of complementary functional assays limit our ability to link transcript-level preservation to functional muscle phenotypes. We also had limited characterization of the cultures so we could not directly relate transcriptional preservation to cell culture quality or traits (ex. fusion index). Future studies incorporating multi-omics integration and advanced culture systems with disease-relevant environmental stimuli may further clarify how transcript-level preservation translates into functional muscle phenotypes.

In summary, gene expression signatures associated with COPD and COPD cachexia in bulk skeletal muscle biopsies were broadly preserved across myoblast, myocyte, and myotube differentiation stages, with myoblast and myotubes most completely recapitulating disease-relevant transcriptional signatures. These results provide a foundation for myoblasts and myotubes as in vitro models of degeneration and repair pathway dysregulation in COPD-cachexia, while myocytes retain select disease-relevant programs reflective of their transitional state. Together, these findings support the use of patient-derived HMDC cultures as tractable systems for testing candidate pathways and interventions implicated in COPD-cachexia.

## ACKNOWLEDGEMENTS

The authors extend sincere gratitude to the UAB Genomics and Sequencing Core, which is supported by Sequencing Core supported by S10OD032422, as well as UAB’s Metabolism Core, which is supported by the UAB Diabetes Research Center (P30DK079626) and the UAB Center for Clinical and Translational Science (UL1TR003096). Additional thanks to Kimberly Estell, Vivian Lin, and Kevin Mitchell in the UAB Lung Health Center biorepository.

## ETHICS STATEMENT

Participants were informed about study procedures and risks before giving written informed consent and starting the study protocol. The study was approved by the University of Alabama at Birmingham Institutional Review Board (IRB-161017009) and conducted in accordance with the ethical standards laid down in the 1964 Declaration of Helsinki and its later amendments.

## AUTHOR CONTRIBUTIONS

Conception and design of research: MLM, ATM, RC, RP, JMW, HBR; Performed experiments: KML, HS, BLS, WM; Drafted manuscript: HS, BLS, MLM; Prepared figures and tables: KML, HS,; Provided feedback on first draft of manuscript: JWC, RP, JMW, HBR, ATM, MLM; Approved final version of manuscript: HS, BLS KML, WM, JWC, RC, RP, JMW, MK, HBR, ATM, MLM

---

**Fig. S1.**
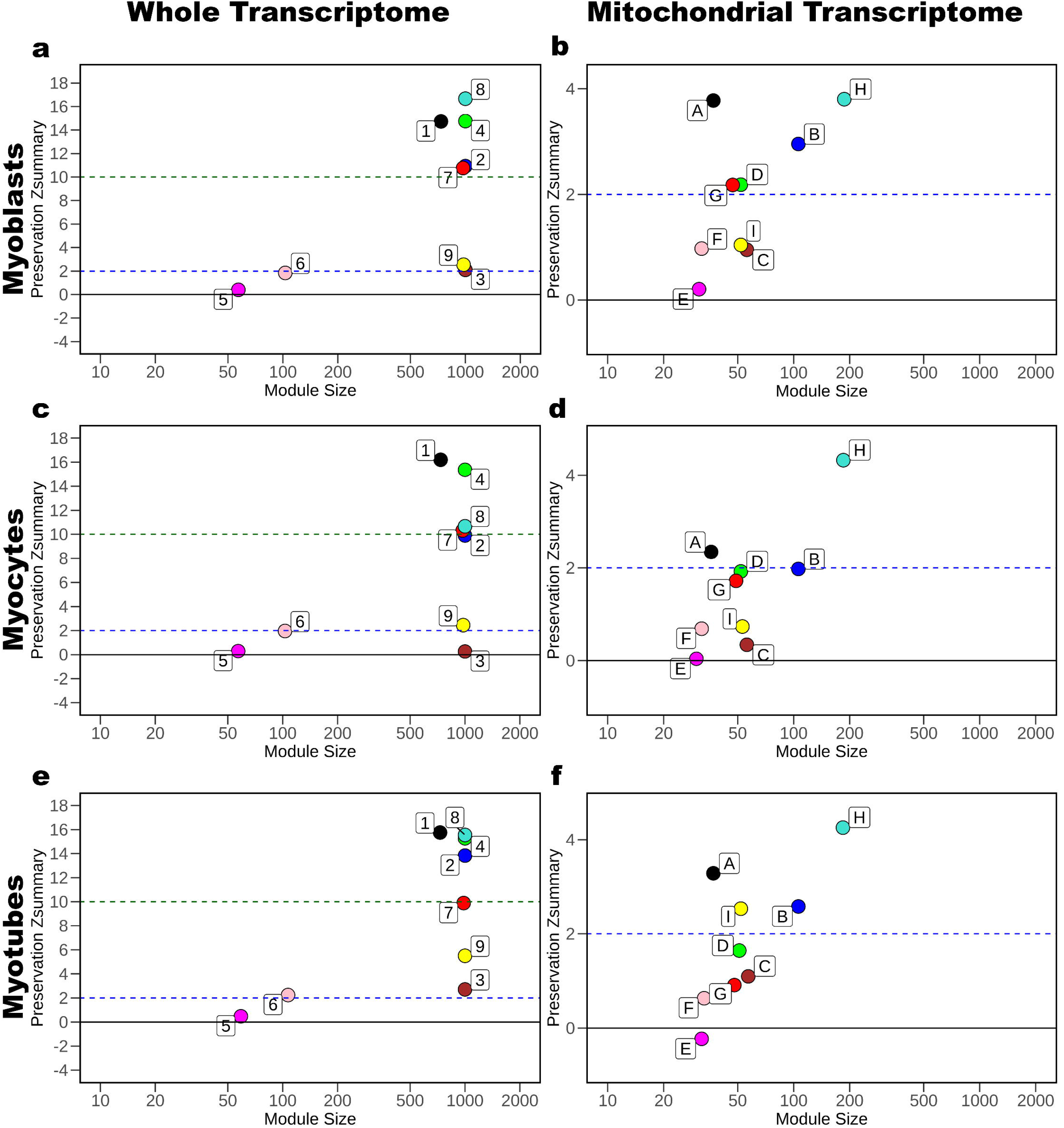
Preservation of Bulk Skeletal Muscle Modules in HMDC states from the whole-and mitochondrial transcriptome networks. Z-Summary scores and module overlap size for mature skeletal muscle network vs (a & b) myoblasts, (c & d) myocytes, and (e & f) myotubes. Plots were created in R using the ggplot2 package.

